# Cross-Species Communication via Fungal Extracellular Vesicles

**DOI:** 10.1101/2025.02.03.636213

**Authors:** Renan E. A. Piraine, Henrique T. Oliveira, Patrick W. Santos, Julia L. Froldi, Bianca T. M. Oliveira, Caroline P. Rezende, Gabriel E. S. Trentin, Lucas F. B. Nogueira, Arnaldo L. Colombo, Arturo Casadevall, Marcio L. Rodrigues, Fausto Almeida

**Affiliations:** Department of Biochemistry and Immunology, Ribeirão Preto Medical School, University of São Paulo, Ribeirão Preto, São Paulo, Brazil; Department of Molecular Biology, Cancer Molecular Research Laboratory (LIMC)/ FAMERP, São José do Rio Preto, São Paulo, 15090-000, Brazil; Special Laboratory of Mycology, University of São Paulo, São Paulo, São Paulo, Brazil; Department of Molecular Microbiology and Immunology, Johns Hopkins School of Public Health, Baltimore, Maryland, United States; Gene Expression Regulation Laboratory, Carlos Chagas Institute, FIOCRUZ, Curitiba, Paraná, Brazil

**Keywords:** virulence factors, pathogenic fungi, intraspecies, interspecies, immunostimulation, fungal communication

## Abstract

Extracellular vesicles (EVs) play crucial roles in fungal communication and host immune modulation, representing potential therapeutic targets for fungal infections. This study investigated the role of fungal EVs in both intra- and interspecies communication, focusing on their effects on virulence and immune responses. Co-incubation experiments were performed using EVs derived from *Candida albicans* and *Candida auris* to assess interactions with *C. albicans* planktonic cells and biofilms, as well as *Cryptococcus neoformans* and *Cryptococcus gattii* EVs interacting with *C. neoformans* cultures. EVs were observed associating with recipient cell surfaces, suggesting subsequent internalization. Functional assays revealed that EV exposure led to increased expression of *Cap59, Lac1, Ure1,* and *Erg11* genes, correlating with reduced antifungal susceptibility in both planktonic and biofilm forms. Additionally, EVs facilitated cross-species communication, enhancing biofilm adhesion and dispersion, which underscores their role in phenotypic modulation. Macrophages stimulated with fungal EVs exhibited receptor-specific gene expression changes, notably the upregulation of *galectin-3*, along with a pro-inflammatory phenotype marked by increased *iNOS* expression and elevated cytokine levels (IL-1β, IL-6, and IL-8). Collectively, these findings underscore a critical role for fungal EVs in interspecies communication, biofilm regulation, and immune modulation, offering valuable insights into fungal pathogenicity mechanisms.

**Importance:** Currently, no vaccines exist to prevent fungal infections, underscoring the need for new therapies. As fungal diseases increase globally, understanding fungal biology is essential to identifying treatment targets. Fungi use EVs to communicate and evade immune responses. EVs mediate cell-cell communication, transporting proteins, polysaccharides, lipids, and nucleic acids – serving as “messages” exchanged within a fungal network. Understanding how these vesicles facilitate communication not only within a single species but also across different fungal species can shed light on their contribution to infection persistence and cross-species adaptability. Moreover, EVs may have a broader role in inter-kingdom communication, influencing how fungi interact with host immune cells. The impact of fungal EVs on human innate immune responses remains a largely underexplored area, with significant gaps in our knowledge. This study aims to examine how fungal EVs affect immune responses and whether their signaling varies across species, potentially revealing new therapeutic targets.

## Introduction

In recent decades, fungi have emerged as significant human pathogens, posing a global threat to human health. This trend highlights the urgent need to address fungal infections, particularly with the rise of multidrug-resistant strains (1). The increase in fungal infections is closely associated with the rising number of immunocompromised individuals, as well as the fungi’s own evolving capabilities, such as enhanced pathogenicity and the ability to colonize new ecological niches (2–4). Fungal infections now affect over 1 billion people annually, with 2.55 million deaths reported worldwide, particularly among immunocompromised individuals (2, 5). Despite the considerable global health burden, the U.S. Food and Drug Administration (FDA) has yet to approve any vaccines for fungal infections (6). In 2022, the World Health Organization published its first ever *fungal priority pathogens list*, which categorizes pathogens into critical, high, and moderate threats. Among the "critical importance" group are *Aspergillus fumigatus*, *Candida albicans*, *Candida auris*, and *Cryptococcus neoformans*. These pathogens represent significant challenges due to their virulence and resistance to treatment. (3, 7).

*Candida albicans,* and more recently *C. auris,* are among the most prevalent *Candida* species associated with candidiasis in humans, causing nosocomial outbreaks with mortality rates exceeding 40% in some studies (1, 8). Effective control strategies, including the use of antifungal therapies and rapid diagnostic tools, are crucial to prevent disease progression. Similarly, cryptococcosis, caused by *C. neoformans* and *C. gattii,* poses a comparable mortality risk due to severe pulmonary and central nervous system infections (1, 9). The ability of both *Candida* and *Cryptococcus* to establish infections is facilitated by several key virulence factors, such as thermotolerance, dimorphism, biofilm formation, capsule production, adhesins, and hydrolytic enzymes, all of which contribute to their pathogenicity and resistance to host defenses (3).

Fungal EVs play a critical role in pathogenesis, often described as “virulence bags” due to their role in transporting a variety of bioactive molecules, including proteins, lipids, nucleic acids, polysaccharides, toxins, and pigments. These components facilitate fungal dissemination and immune evasion within the host environment (10), functioning as carriers for molecules destined for the extracellular space (11). EVs are spherical structures enclosed by a bilayer membrane, categorized into small (<200 nm) and large (>200 nm) vesicles, following MISEV2023 (*Minimal information for studies of extracellular vesicles*) guidelines (12). Our group has shown that fungal EVs play a key role in facilitating communication among fungal cells within the same species (intraspecies interactions) (13). Additionally, these EVs also act as immunomodulators in host-pathogen interactions, as evidenced by *in vitro* studies involving immune cells (14–17).

Fungi can internalize EVs, triggering a variety of responses, including changes in gene expression. These responses are critical for fungal pathogens, influencing key processes such as antifungal tolerance, virulence, morphogenesis, and growth (13, 18). By coordinating fungal communities, EVs enhance their survival and adaptability within the host environment (19). EVs play a significant role in *C. albicans* biofilm formation and the yeast-to-hypha transition (13, 20, 21). However, further studies are needed to elucidate the roles of EVs in cross-species signaling and their impact on fungal biology during active infections.

Emerging studies are also investigating interactions between different *Candida* species mediated by EVs, prompted by epidemiological data suggesting that multispecies infections are becoming more common (22). In *Cryptococcus* species, EVs regulate key virulence mechanisms and enable long-distance signaling (23), with research on this topic spanning over the past 15 years (24, 25). Beyond their role in cell wall dynamics, cryptococcal EVs are critical for capsule biogenesis and maintenance, as they transport the major capsule component, *glucuronoxylomannan* (GXM), through a trans-cell wall export mechanism (11). Although coinfections involving different *Cryptococcus* species in humans are considered rare (26, 27), studying EV-mediated interspecies communication provides valuable insights into the conservation of cellular communication mechanisms across fungal species.

In addition to their role in cellular communication, EVs are potent stimulators of host immunity (28). Most research on the immunostimulatory effects of fungal EVs has been conducted *in vitro* using immune cells, primarily focusing on species from the *Candida* and *Cryptococcus* genera (28, 29). EVs from *C. albicans* were implicated in modulating both innate and adaptive immune responses, including the induction of nitric oxide (NO) production and the regulation of inflammatory response (30, 31). Key virulence factors in *C. albicans* EVs that contribute to host immune activation include phosphatidylserine, glucosylceramide, and small RNAs (28, 31). In *Cryptococcus*, EVs also carry important virulence factors such as glucosylceramide and small RNAs. However, two other prominent factors, GXM and melanin, are known for their cytotoxic effects and ability to suppress phagocytosis, respectively (28, 32). While some studies report an anti-inflammatory response, characterized by increased IL-10 and TGF-β (Transforming Growth Factor β) in macrophages stimulated with *C. neoformans* EVs (29, 33), other research has observed pro-inflammatory responses following stimulation with EVs from *C. neoformans* and *C. gatti* (32, 34).

In this study, we aimed to explore the role of fungal EVs in communication and immune modulation, isolating and characterizing EVs from *C. albicans, C. auris, C. neoformans,* and *C. gattii.* We investigated intra- and interspecies communication by assessing EVs uptake and impact on planktonic cells, biofilm formation, antifungal tolerance, and the transfer of capsular material. Additionally, we examined the immunomodulatory activity of these EVs on THP-1 human macrophage-like cells. This analysis focused on internalization, potential cytotoxic effects, the production of pro- and anti-inflammatory cytokines, and the mRNA expression of immune response-associated genes.

## Results

### Fungal EVs as mediators in intra- and inter-species communication

The detailed characterization of EVs structure, cargo, and biological significance began around 2007 with the study of EVs from *C. neoformans* (29). Understanding the characteristics of EVs may reveal species-specific attributes that are critical for their function. Figure 1 (A-D) presents the results of Nanoparticle Tracking Analysis (NTA) and Transmission Electron Microscopy (TEM) for EVs isolated from cultures of *C. albicans* ATCC64548, *C. auris* clinical isolate 470/2015, *C. neoformans* H99, and *C. gattii* R265. Characterization of these EVs revealed a heterogeneous population in all samples, ranging from small EVs (sEVs) with a size of < 200 nm to large EVs (lEVs) with > 200 nm (summarized in Fig. 1E). The yield of EVs recovered after the isolation protocol (centrifugation, filtration, and ultracentrifugation) was quantified by the EV-to-fungal cell ratio, a method suggested for evaluating and comparing EV isolation from solid media-based cultures. Zeta potential, an important indicator of surface charge and stability, was also measured using Dynamic Light Scattering (DLS). The zeta potential of fungal EVs ranged from -8 to -15 mV, indicating a stable colloidal suspension under physiological conditions (Fig. 1F).

**Figure 1:**
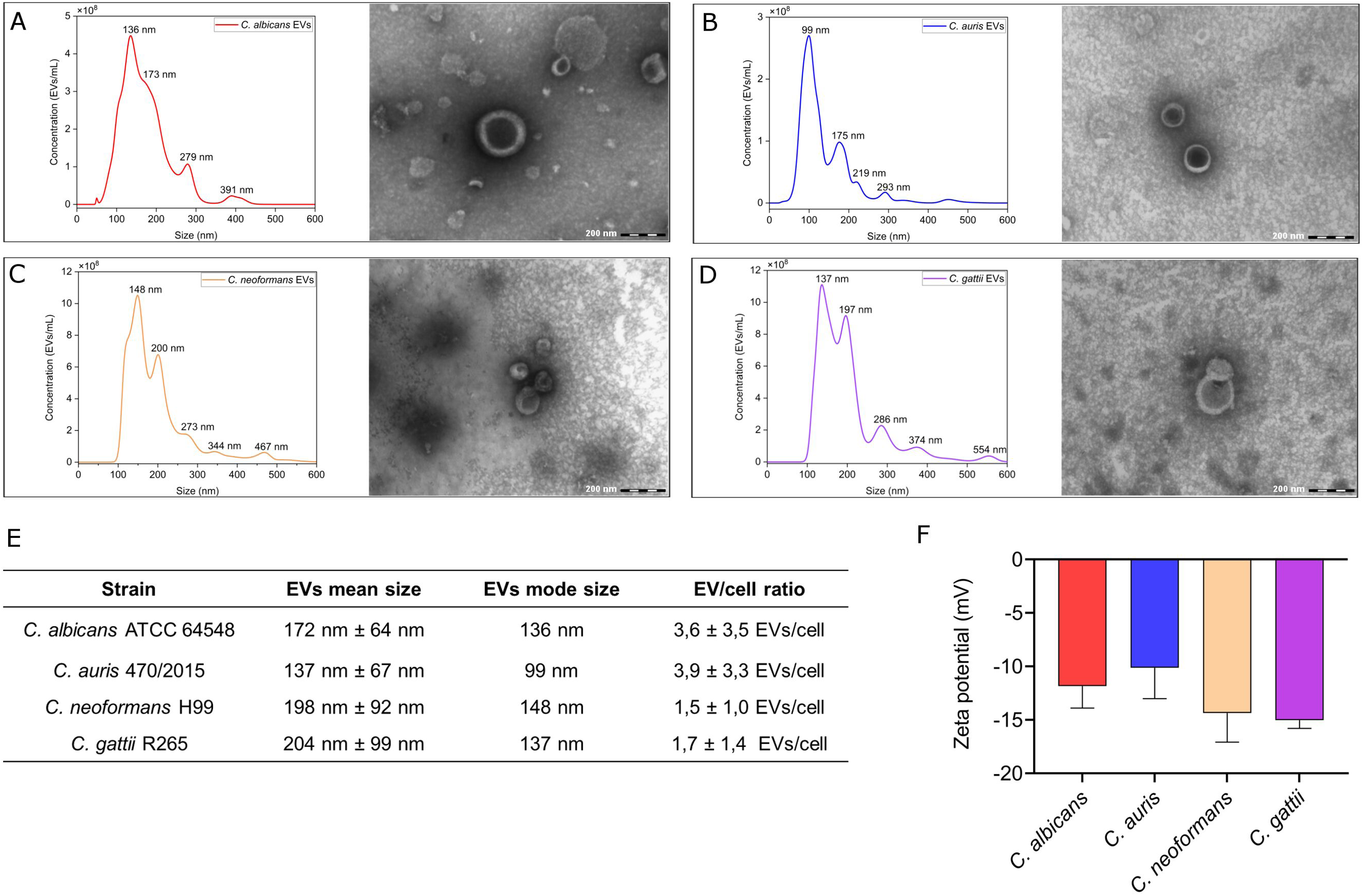
Characterization of fungal extracellular vesicles (EVs). EVs isolated from *C. albicans* ATCC 64548 (A), *C. auris* 470/2015 (B), *C. neoformans* H99 (C), and *C. gattii* R265 (D) were analyzed by Nanoparticle Tracking Analysis (NTA). The data obtained from NTA is summarized in panel (E). Zeta potential measurements of EVs were performed using Dynamic Light Scattering and are shown in panel (F). Statistical analysis was conducted using One-way ANOVA in GraphPad Prism 9.0. Error bars represent mean ± standard deviation.

Our group previously demonstrated the role of fungal EVs in intraspecies communication (13). In this study, we explore the potential for EVs to serve as vehicles for interspecies communication, focusing on their ability to be “sent” and “received” as messages between fungi of different species. Using labeled EVs and fluorescence microscopy, we observed the recognition and possible internalization of EVs from different species. Specifically, we detected interactions between *C. albicans* cells and *C. auris* EVs, *C. gattii* cells and *C. neoformans* EVs, and an intergeneric interaction between *C. auris* cells and *C. neoformans* EVs (Fig. 2A). Additionally, scanning electron microscopy (SEM) revealed an increase in vesicle-like structures adhered to the surface of recipient cells after incubations with EVs isolated from other species, supporting the hypothesis that EVs interact with the fungal cell wall and/or membranes before internalization (Fig 2B and C).

**Figure 2:**
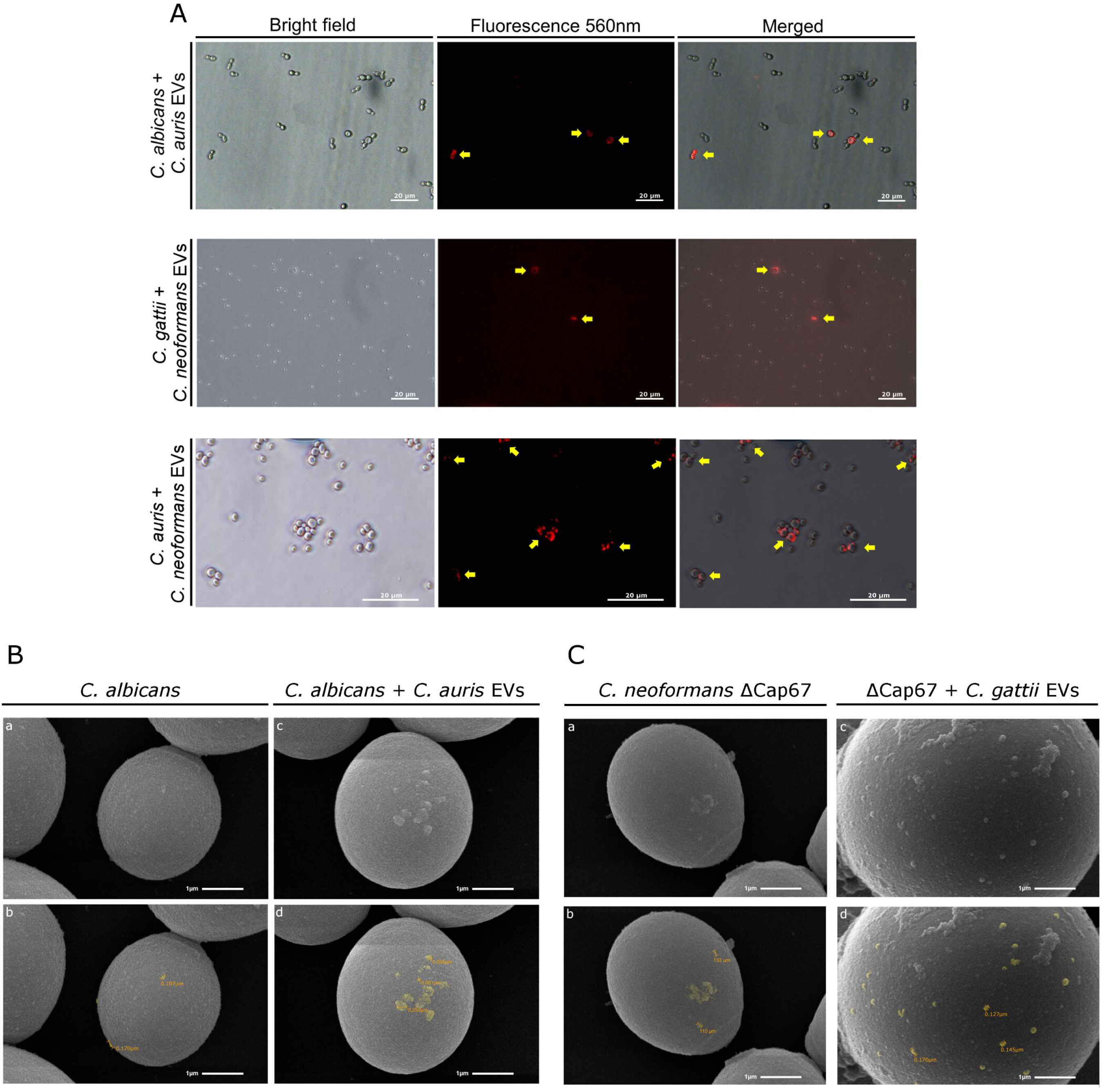
Interspecies cell surface association and internalization of fungal EVs. (A) Fluorescence microscopy images suggest that fungal cells are capable of recognizing, interacting with, and possibly internalizing EVs produced by different species and genera. Yellow arrows indicate cells that are internalizing EVs, at 400 × magnification. SEM images show the association of fungal EVs (indicated in yellow) with the cell wall after co-incubation with *C. albicans* (A) and *C. neoformans* ΔCap67 (B), demonstrating interspecies interactions between EVs and fungal cells. Images “b” and “d” in both experiments (for *Candida* spp. and *Cryptococcus* spp.) were manually colored using GIMP2® software to correspond to images “a” and “c”. Magnification of 20,000x.

Extracellular vesicles are frequently referred to as “virulence bags,” as they transport proteins and other biomolecules between cells (3). In our study, we explored the biological impact of EVs in interspecies communication, using *C. neoformans* ΔCap67 (an acapsular mutant) as a model to better understand the role of EVs in the transmission of virulence factors. We chose the *C. neoformans* ΔCap67 strain to avoid the capsule when exploring EV delivery, allowing for a more detailed examination of the cell surface without interference from capsular components that could hinder the visualization of EV incorporation. This approach was particularly valuable for investigating the transfer of GXM and other associated molecules from EVs derived from wild-type strains (H99 or R265), to determine whether the transfer of EVs cargo is sufficient to restore the mutant phenotype. While *C. neoformans* var. *grubii* H99 is classified as serotype A, the ΔCap67 mutant is a serotype D strain derived from strain B3501 (*C. neoformans* var. *neoformans*) (35). ΔCap67 is also referred to as strain B4131 in the literature, and, as well as the ΔCap59 strain (an acapsular mutant derived from H99), its phenotype can be restored by *Cap59* gene complementation (both strains show mutations in *Cap59*) (35, 36).

We firstly examined the effect of adding EVs to planktonic cells to assess their impact on virulence factor expression at the molecular level. Specifically, we selected three genes - *Cap59*, *Lac1*, and *Ure1* – and measured their transcription by qPCR. Our results showed increased expression of these genes when ΔCap67 mutant cells were incubated with EVs from *C. neoformans* and *C. gattii* (Fig. 3A-C). This response was observed in both intra- and interspecies interaction. Even *C. neoformans* ΔCap67 EVs (Fig. S1A – Supplementary Information for *C. neoformans* ΔCap67 NTA data) induced a modest increase in mRNA expression of these genes, albeit at lower levels. Overall, the addition of EVs led to an upregulation of all tested genes, suggesting that EVs directly contribute to the induction of virulence factor production.

**Figure 3:**
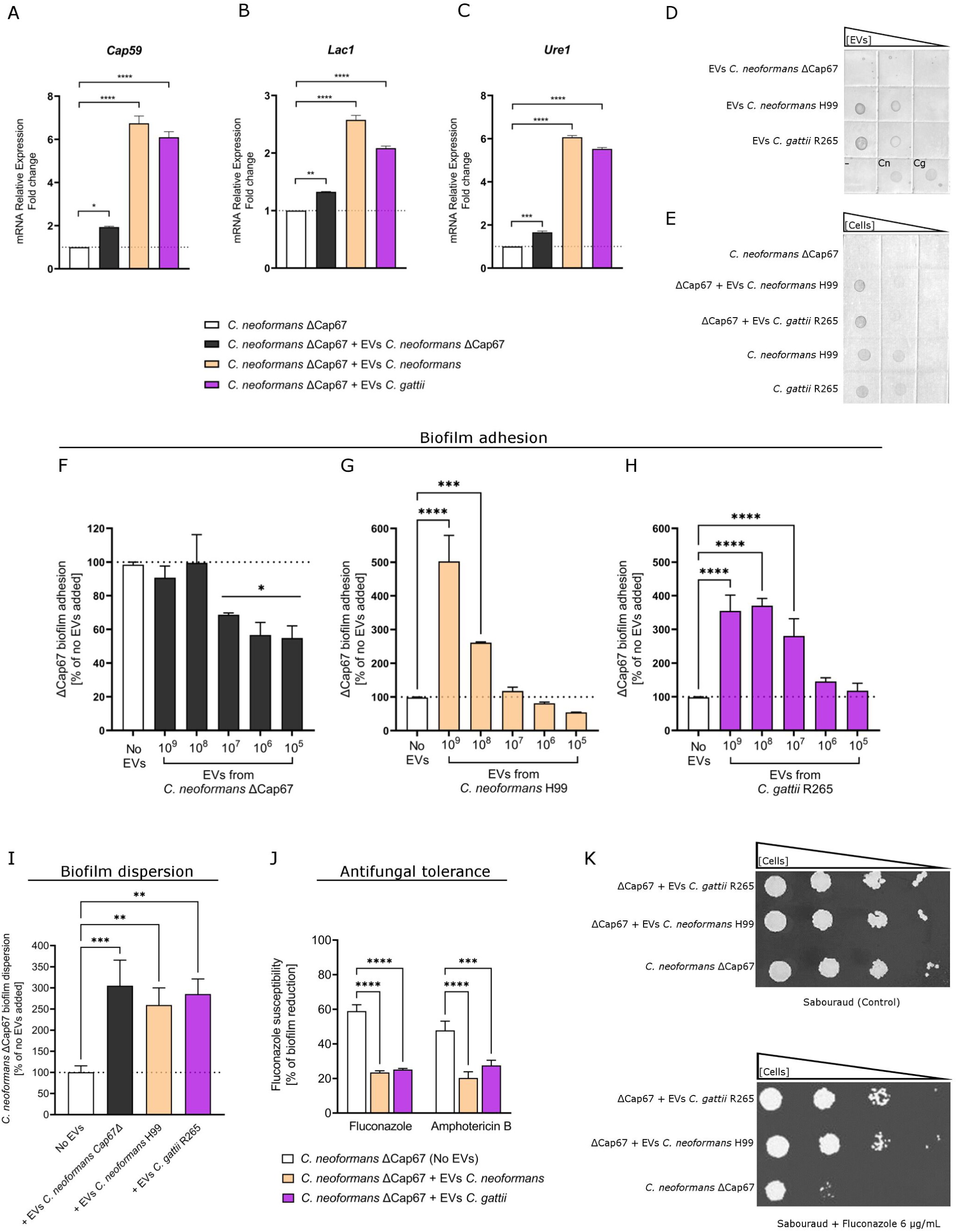
Biological impact of EVs derived from *C. neoformans* and *C. gattii* on planktonic cells and biofilms. Planktonic cells of *C. neoformans* ΔCap67 were incubated with EVs isolated from *C. neoformans* H99 and *C. gattii* R265. mRNA relative expression levels of *Cap59 (*A)*, Lac1* (B), and *Ure1* (C) were measured by qPCR and compared to the control (no EV treatment). The dotted lines represent the basal expression in *C. neoformans* ΔCap67 cultures. (D) A dot blot was used to detect the presence of the capsular content GXM in fungal EVs. BSA 5% served as a negative control (-), while *Cn*: *C. neoformans* H99, and *Cg*: *C. gattii* R265 cultures were used as positive controls. (E) Dot blot analysis of ΔCap67 cultures incubated with EVs from *C. neoformans* H99 and *C. gattii* R265, showing a positive reaction with the anti-GXM monoclonal antibody. (F-H) EVs from *C. neoformans* ΔCap67, *C. neoformans* H99, and *C. gattii* R265, at concentrations ranging from 10^4^ to 10^8^ EVs/well, were added to biofilms of *C. neoformans* ΔCap67. Biofilm adhesion was evaluated at each EV concentration, and the concentration with the most pronounced biological effect was selected for further analysis of biofilm dispersion (I) and susceptibility to fluconazole and amphotericin B (J). The dotted lines represent the biofilm formed by non-treated *C. neoformans* ΔCap67 (no EVs added). Statistical analysis was performed using One-way ANOVA in GraphPad Prism 9.0, with significant differences indicated by the “*” symbol: *p < 0.05, **p < 0.01, ***p < 0.001, and ****p < 0.0001. Error bars represent means ± standard deviation. (K) Spot tests were conducted with dilutions of *C. neoformans* ΔCap67 cultures, with or without EVs (control), to assess the effect on antifungal tolerance in planktonic cells.

Next, we confirmed the presence of GXM in the EVs of *C. neoformans* H99 and *C. gattii* R265, and its absence in *ΔCap67* mutant EVs, using a dot blot assay (Fig. 3D). Co-incubation of EVs from wild-type strains (H99 and R265) with ΔCap67 mutant cells revealed that GXM was incorporated into the mutant’s surface, indicating the uptake of a significant amount of the polysaccharide (Fig. 3E). SEM images (Fig. 2C) further demonstrated that the acapsular mutant did not fully acquire a capsular structure after 24 h of co-incubation, in contrast to the complete phenotype reversal seen in the wild-type strain. This was further confirmed by India ink staining, where no significant differences were observed between cells that received EVs and the controls (no EVs added) (Fig. S2 - Supplementary Information).

The characteristics of *C. neoformans* ΔCap67 biofilms were also altered after exposure to EVs. While no significant differences in adhesion were observed when biofilms were incubated with ΔCap67 EVs (compared to biofilms with no EVs added), the highest concentrations of EVs from *C. neoformans* H99 and *C gattii* R265 (10^9^ and 10^8^ EVs/well, respectively) resulted in a significant increase in the number of adhered cells during the adhesion phase (Fig. 3F-H). The highest concentration of EVs (10^9^ EVs/well) was also tested in the biofilm dispersion assay, where it was shown that EVs influence the dispersion phase. However, no statistically significant differences were observed in the levels of dispersion between EVs from the same or different strains (Fig. 3I). Additionally, the impact of EVs on antifungals susceptibility was assessed, revealing that biofilms incubated with EVs from the wild-type strains of *C. neoformans* and *C. gattii* exhibited increased tolerance to antifungals (Fig. 3J). The effect on virulence factors was further confirmed by a spot test, where planktonic cells co-incubated with EVs from *C. neoformans* and *C. gattii* showed reduced antifungal susceptibility on agar plates containing various concentrations of fluconazole (Fig. 3J and Fig. S3 - Supplementary Information).

The biological impact of fungal EVs on interspecies communication was further demonstrated in both planktonic cells and biofilms of *Candida* yeasts. In planktonic *C. albicans* cells, EVs isolated from both *C. albicans* and *C. auris* induced an increase in mRNA expression of *Erg11*, a gene that is up-regulated in azole-resistant isolates. EVs isolated from *C. auris* cultured in fluconazole-containing plates (see Fig. S1B – Supplementary Information for *C. auris* NTA result) also stimulated elevated *Erg11* expression in planktonic *C. albicans*. Notably, qPCR analysis revealed that the increase in *Erg11* expression was approximately 2-fold higher than that observed with EVs from *C. albicans* or untreated *C. auris* EVs (Fig. 4A). Additionally, spot tests demonstrated a difference in fluconazole susceptibility between planktonic *C. albicans* cells incubated with EVs from *C. auris*, in which at high fluconazole concentrations, *C. albicans* cells exposed to *C. auris* EVs exhibited a more tolerant phenotype (Fig. 4B).

**Figure 4:**
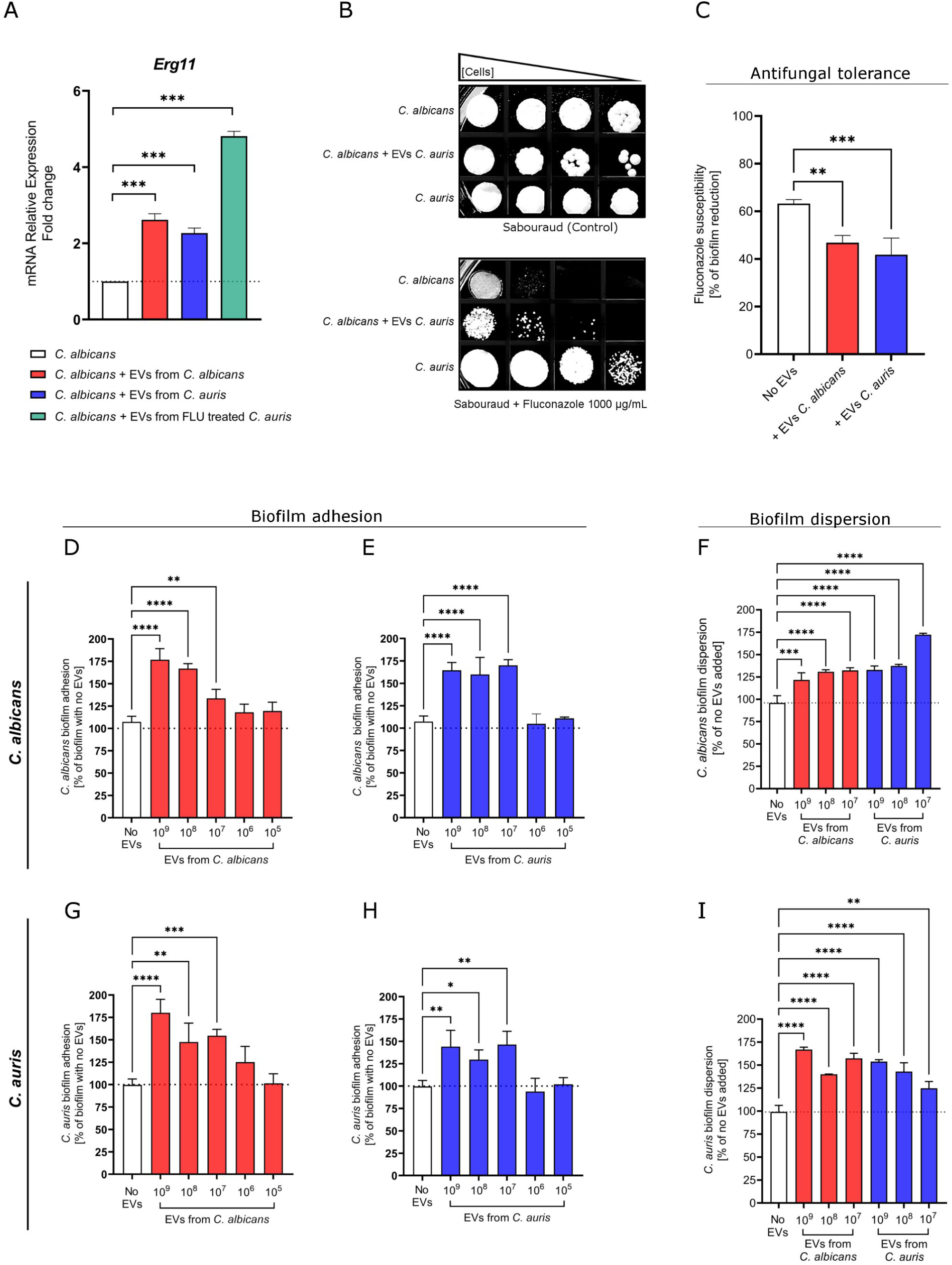
Biological impact of EVs from *C. albicans* and *C. auris* on planktonic cells and biofilms. (A) Planktonic *C. albicans* cells were incubated with EVs isolated from *C. albicans*, *C. auris*, or *C. auris* treated with fluconazole. *Erg11* mRNA relative expression was measured by qPCR and compared to the control (no EV treatment). The dotted line represents the basal expression in *C. albicans* cultures. (B) Spot tests were conducted with dilutions of *C. albicans* cultures that were treated with or without EVs (control strain: *C. albicans* and *C. auris*) on Sabouraud agar plates with or without fluconazole. (C) Fluconazole susceptibility in *C. albicans* (a fluconazole-susceptible strain) biofilms was tested following the addition of *C. albicans* EVs or *C. auris* EVs (10^9^ EVs/mL). (D, E, and F) Different concentrations of EVs from *C. albicans* and *C. auris* were added on adhesion and dispersion phase of *C. albicans* biofilms. (G, H, and I) Different concentrations of EVs from *C. albicans* and *C. auris* were added on adhesion and dispersion phase of *C. auris* biofilms. Biofilm dispersion was assessed after adding EVs to pre-formed biofilms of both *Candida* species. For all biofilm assays, the dotted lines represent results for biofilms with no EV addition. Statistical analysis was performed using One-way ANOVA in GraphPad Prism 9.0, with significant differences indicated by the “*” symbol: *p < 0.05, **p < 0.01, ***p < 0.001, and ****p < 0.0001. Error bars represent means ± standard deviation.

The cross-communication between *C. albicans* and *C. auris* through EVs was evaluated using assays of biofilm adhesion, dispersion, and fluconazole tolerance. As seen in planktonic assays, exposure to EVs also altered azole tolerance in *C. albicans*. In *C. albicans* biofilms, fluconazole treatment resulted in an approximately 62% reduction in cell viability of biofilm mass (control). However, biofilms treated with *C. albicans* EVs showed a reduction of approximately 47%, and those treated with *C. auris* EVs exhibited a reduction of 42%, suggesting a decreased susceptibility to fluconazole in the presence of EVs (Fig. 4C). In biofilms of both *C. albicans* and *C. auris*, fungal cell/EV ratios ranging from 1:10 to 1:1000 (10^7^ to 10^9^ EVs/well) resulted in significantly increased biofilm adhesion compared to the control (no EVs added). No differences were observed when EV concentrations were < 10^6^ EVs/well (Fig. 4D, E, G, H). The EV concentrations that enhanced biofilm adhesion were selected for further investigation of their impact on biofilm dispersion. In these assays, biofilms treated with EVs, whether from *C. albicans* or *C. auris*, showed higher levels of dispersed cells compared to the control (Fig. 4 F and I), indicating that both intraspecies and interspecies communication promoted biofilm dispersal.

### Fungal EVs as molecules with immunomodulatory properties – the inter-kingdom communication

One of the critical factors in host-pathogen interactions during infection is communication via EVs (34). Fungal EVs play a pivotal role in fungal pathogenicity, as they transfer virulence factors with immunomodulatory properties. These factors are recognized as pathogen-associated molecular patterns (PAMPs), thereby activating the innate immune response (3, 35). Similar to studies conducted with fungal cells, we labeled fungal EVs with a fluorescent dye to observe their incorporation or internalization by immune system cells – specifically, human macrophage-like cell cultures (THP-1 cells). Through fluorescence microscopy, we detected labeled EVs overlapping with regions where THP-1 cells were located (Fig. 5A), suggesting an association with the macrophage-like cell surface and potential internalization. Since the cytotoxic effects of fungal EVs vary depending on the study and experimental conditions, we further investigated their impact on cellular properties during co-incubation of THP-1 cells with EVs from *Cryptococcus* spp. and *Candida* spp. The assay revealed no significant differences in human macrophage-like cells cultured *in vitro* concerning cell size, viability, or concentration following exposure to fungal EVs (Fig. 5B-D).

**Figure 5:**
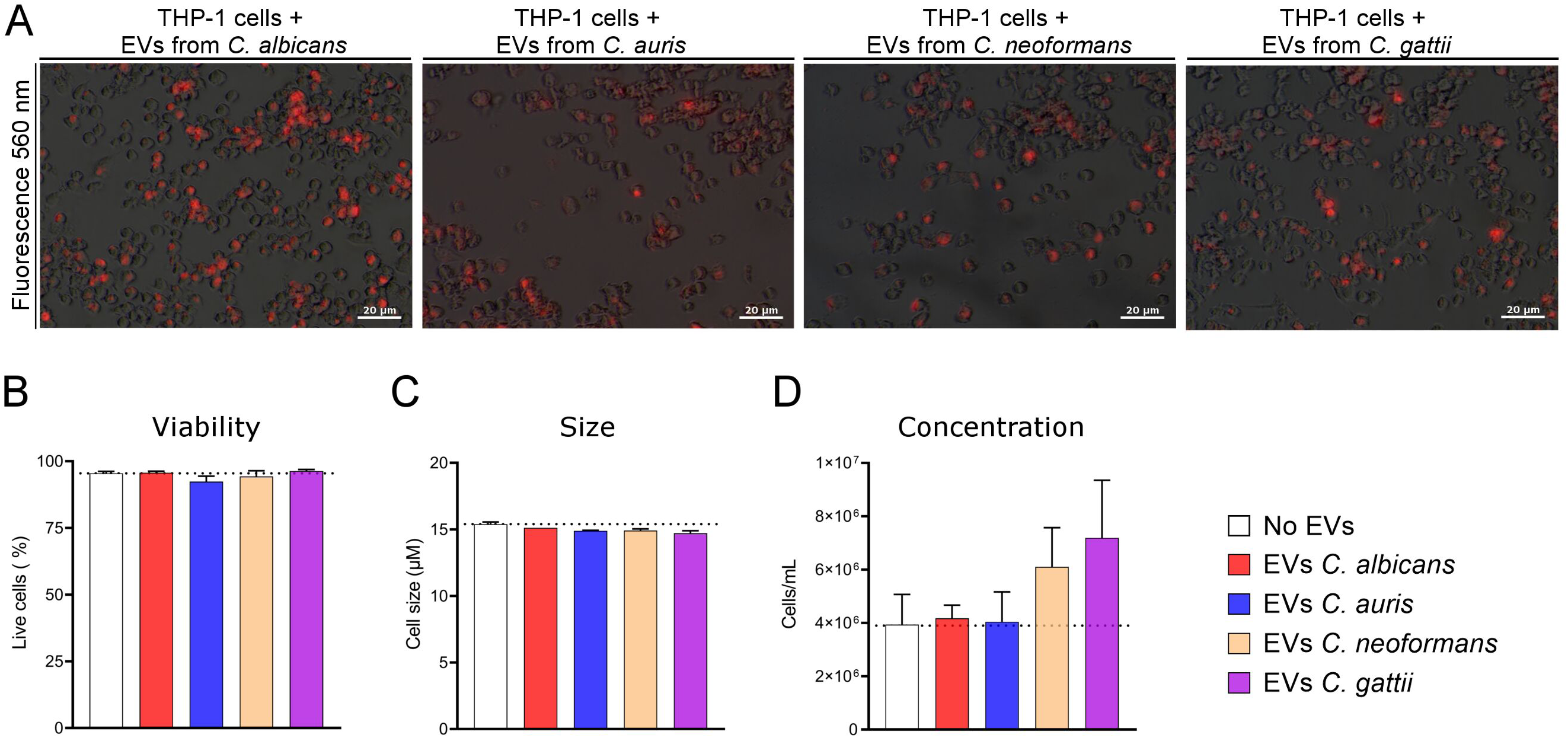
Interaction between fungal EVs and THP-1 macrophages, and its impact on cellular properties. (A) The internalization of EVs by THP-1 cells was observed through fluorescence microscopy, demonstrating successful recognition and interaction with human macrophages. Micrographs from bright field and under fluorescence (560 nm) were merged using GIMP® 2 software. The biological effects on THP-1 cells following EV interaction were assessed by measuring the number of live cells (B), cell size (C), and cell concentration (D). Dotted lines indicated the data observed for untreated cells (control).

The stimulation of THP-1 macrophage-like cells by fungal EVs was analyzed using qPCR, targeting the mRNA transcription of polarization markers, receptors, and proteins involved in immune response pathways associated with combating pathogenic fungi (Fig. 6A-M). All fungal EVs induced an up-regulation of *iNOS* (nitric oxide synthase) and either a basal or downregulated expression of *ARG1* (arginase 1) (p < 0.05). This response suggests the polarization of macrophages toward the M1 phenotype.

**Figure 6:**
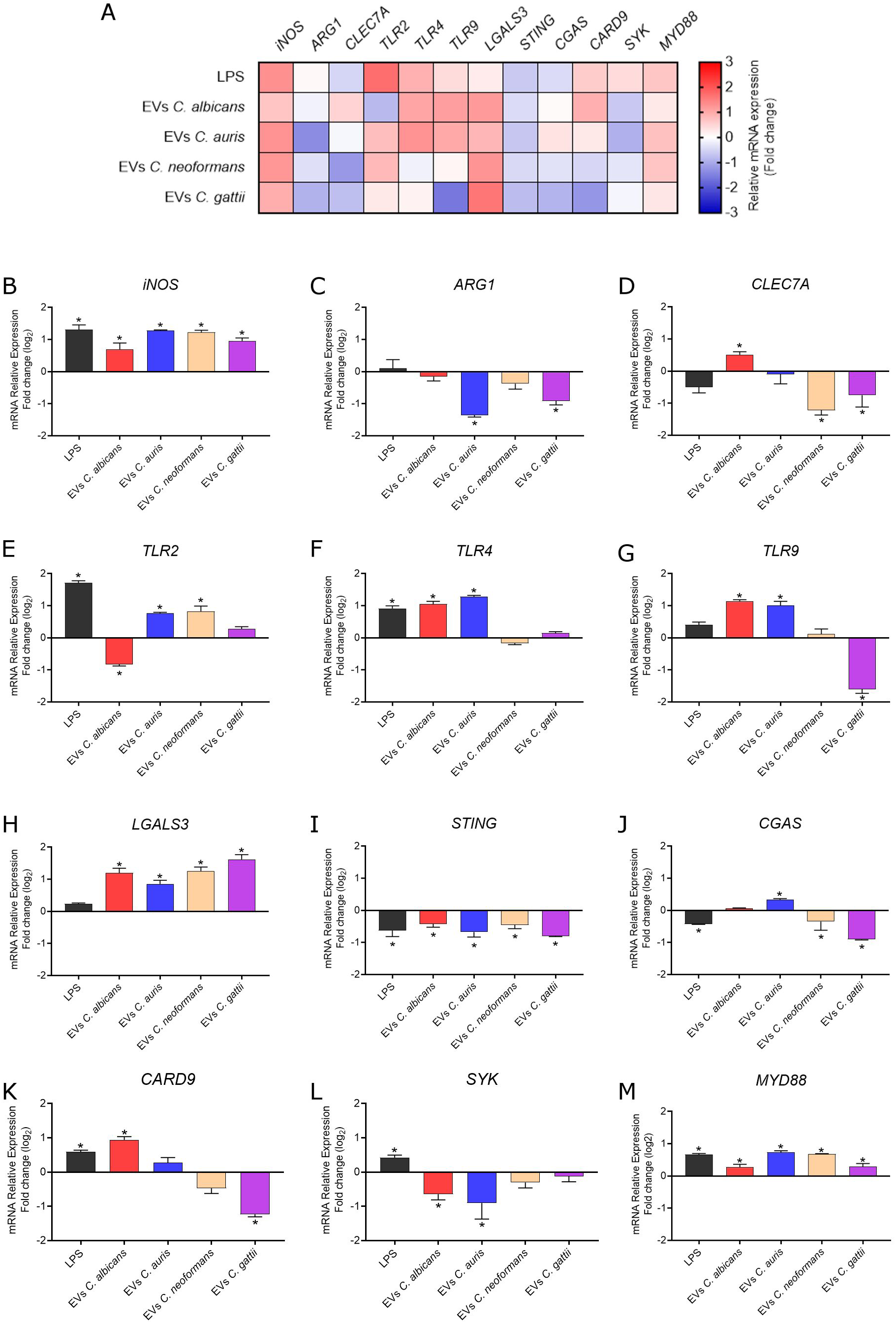
Relative mRNA expression of polarization markers, receptors, and signaling pathway proteins in THP-1 cells stimulated with fungal EVs. This figure presents the relative expression levels of macrophage polarization markers (*iNOS*, *ARG1*), receptors (*TLR2*, *TLR4*, *TLR9*, *CLEC7A*, *LGALS3*), and proteins involved in key immune signaling cascades (*STING*, *CGAS*, *CARD9*, *SYK*, *MYD88*). qPCR results were normalized to the housekeeping β-actin gene (*ACT1*) and calculated using the 2-ΔΔCT method. The heatmap presented in (A) summarizes the results observed for relative mRNA transcription. Fold change for relative mRNA transcription is presented in the graphs (B-M) as log_2_, to facilitate up- and downregulation comparisons.

*Candida* spp. EVs stimulated human macrophages to upregulate the mRNA expression of *TLR4* and *TLR9*. Among these, only *C. albicans* induced significant levels (p < 0.05) of *CLEC7A* expression, while *C. auris* specifically stimulated *TLR2*. In contrast, *Cryptococcus* EVs generally showed either basal mRNA expression (not statistically different) or downregulation of the evaluated receptors. For instance, *CLEC7A* was downregulated in response to EVs from both cryptococcal species, and *TLR9* was downregulated following stimulation with EVs from *C. gattii*. Notably, *TLR2* was the only receptor upregulated by *Cryptococcus* EVs, with its expression significantly enhanced in response to *C. neoformans* EVs. Additionally, *LGALS3*, the gene encoding galectin-3 – a protein that binds beta-galactosides and exhibits antimicrobial activity – was consistently up-regulated in THP-1 cells across all EVs tested.

The immunomodulatory effects of fungal EVs exhibited pattern of *STING* downregulation. Specifically, *Candida* EVs downregulated *SYK* expression, while *Cryptococcus* EVs let to the expression of *CGAS*. Regarding *CARD9*, EVs from *C. neoformans* caused a significant downregulation of its expression, whereas incubation with *C. albicans* EVs resulted in increased mRNA expression in THP-1 cell cultures. *MYD88*, a gene encoding an adapter protein widely studied in immune responses, was upregulated in response to all fungal EV stimuli, with the most pronounced effects observed for EVs derived from *C. auris* and *C. neoformans*.

The quantification of pro- and anti-inflammatory cytokines in the supernatant of macrophage cultures performed by ELISA, allowed the characterization of the immune response profile mediated in THP-1 cells stimulated by fungal EVs. Pro-inflammatory cytokines IL-1β, IL-6, and IL-8 were produced at significantly higher levels compared to the control (non-stimulated cells, medium only) when fungal EVs were present in the medium. However, TNF-α levels, while higher than the negative control, were lower compared to the other pro-inflammatory cytokines (Fig. 7A, C, D, F). Analyzing the response by fungal genera, THP-1 cells stimulated with *C. albicans* EVs produced more IL-6 than those stimulated with *C. auris* EVs, whereas *C. auris* EVs induced higher levels of IL-1β and TNF-α. In the *Cryptococcus* group, EVs from *C. neoformans* stimulated higher levels of IL-1β and IL-8 compared to EVs from *C. gatti*. Regarding the anti-inflammatory cytokines, IL-4 levels remained below the control for all EV stimuli, with very low concentrations being detected. Conversely, IL-10 was produced at levels significantly higher than in control cells (Fig. 7B, E). Taken together, these findings suggest a predominance of a pro-inflammatory response mediated by THP-1 cells in the presence of fungal EVs.

**Figure 7:**
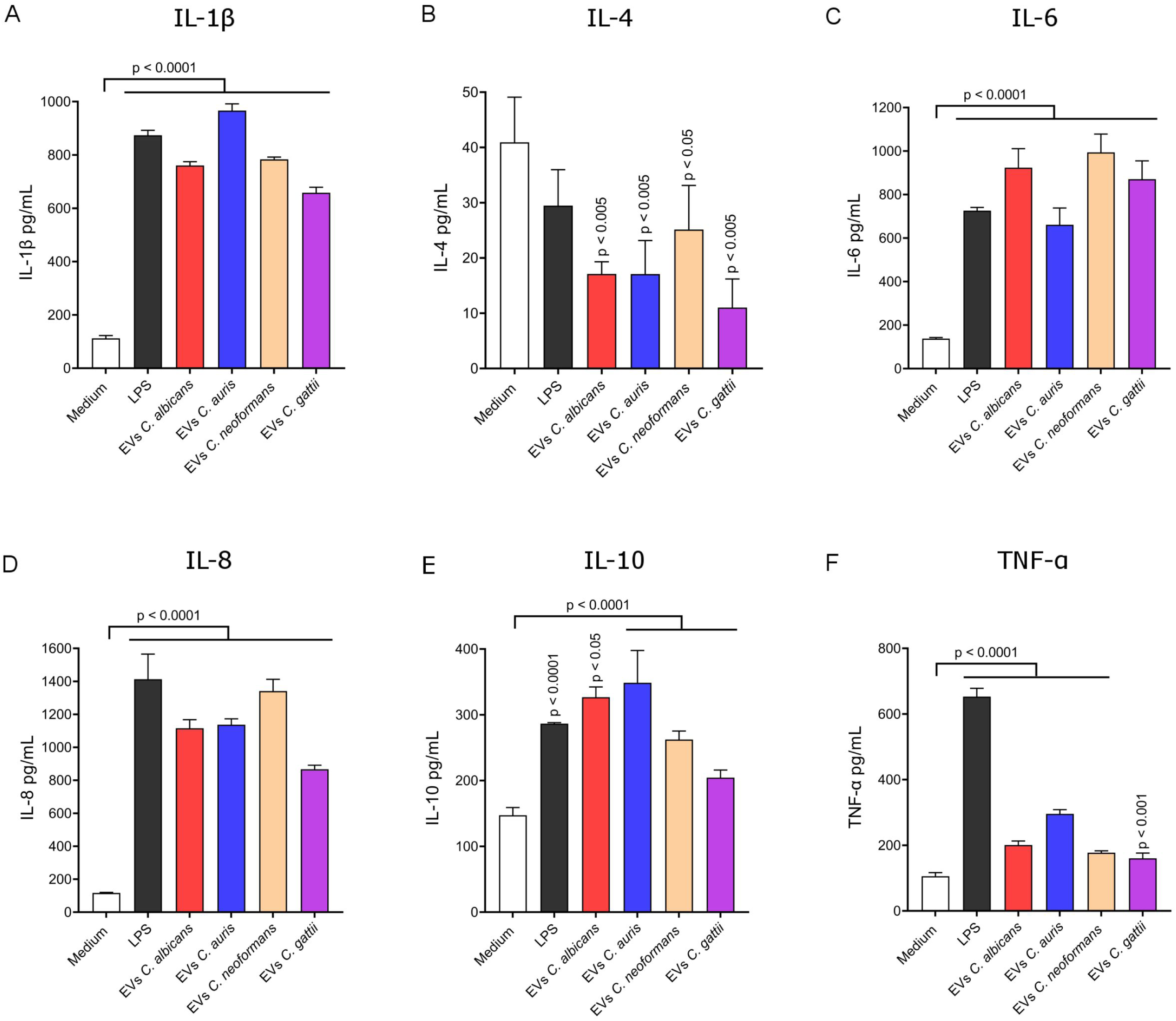
Cytokine production by THP-1 macrophages following stimulation with fungal EVs. THP-1 cells were cultured either with medium only (control) or with EVs derived from *C. albicans*, *C. auris*, *C. neoformans*, and *C. gattii*. The production of IL-1β (A), IL-4 (B), IL-6 (C), L-8 (D), IL-10 (E), and TNFα (F) was quantified by ELISA. Statistical analysis was performed using GraphPad Prism 9.0 with a One-way ANOVA test. Significant differences are indicated by *p*-values above the bars, compared to the control (medium only). Error bars represent the means ± standard deviation.

## Discussion

*Cross-talk* can be defined as an exchange of signals between communicating organisms even over a distance, functioning as a shared “language” based on extracellular signals. These signals are interpreted to coordinate critical processes, such as the survival of pathogenic fungi during infection establishment (37). In fungal communication, essential biological functions are influenced by the exchange of these “messages”, including mating, growth, morphological transitions, and virulence (38). Cross-talk in fungi is partially mediated by EVs - nano-sized structures with bilayer membranes that carry proteins, lipids, glycans, pigments, and nucleic acids (11). Our group recently demonstrated the role of EVs from human pathogenic fungi in intraspecies cellular communication, revealing at the molecular level their impact on gene expression and fungal proliferation control (13). However, this fungal network coordination mechanism does not appear to be limited to the same fungal strain or species. Although still an emerging field, growing evidence supports the existence of EV-mediated interspecies communication, showing significant biological effects, such as biofilm formation and drug-resistance phenotypes, following cross-species interaction (22).

Extracellular vesicles were successfully isolated from cultures of *C. albicans, C. auris, C. neoformans,* and *C. gattii* using the protocol described in Reis et al. (2019) (39). The structure and size of fungal EVs were confirmed through NTA and TEM, corroborating findings from several studies (39–42). These studies consistently described fungal EVs as a heterogeneous population of round-shaped structures, predominantly around 200 nm in size. Regarding the EV-to-cell ratio, our findings align with observations from other research groups (43), which reported up to 10 EVs per cell in similar isolation protocols.

Zeta potential, a measure of particle surface charge, is a key determinant in the stability and functionality of EVs (44). Under physiological conditions, negatively charged particles, including fungal EVs, exhibit enhanced internalization by phagocytic cells such as macrophages, while positively charged particle tend to be preferentially internalized by non-phagocytic cells (44, 45). In this study, fungal EVs consistently exhibited negative zeta potential values, indicating stable colloidal suspensions with reduced likelihood of aggregation. This characteristic support their stability in extracellular environments and suggest increased recognition and uptake by innate immune cells. Similar findings in nanoparticle systems have shown that negative zeta potential reduces protein corona formation, prolongs circulation time, and improves delivery efficiency to target tissues (46). The stability and charge properties of EVs may be leveraged for therapeutic delivery, enhancing biodistribution and targeting infected or inflamed tissues with elevated phagocytic activity (47). This preferential interaction could be further exploited by engineering fungal EVs to carry antifungal compounds or immune modulators, optimizing their delivery to target sites.

It has been demonstrated that vesicular transport in fungi occurs bidirectionally across the fungal cell wall, allowing the export of EVs to the external environment as well as the internalization into intracellular compartments after binding to fungal surface receptors (18, 19). Some studies suggest key processes that may be necessary for EV uptake, including the passage of EVs through cell wall pores approximately 5 nm in size (or up to 400 nm under stress conditions), enzymatic modifications of the cell wall (mediated by enzymes carried by EVs), and transit through specific cellular channels (18, 48). Our group previously reported the uptake and association of radiolabeled EVs by cells of different fungal species, such as *C. albicans, Paracoccidioides brasiliensis,* and *A. fumigatus* (13). In the present study, we demonstrate similar events in *C. albicans*, *C. auris*, *C. neoformans*, and *C. gattii* using a fluorescent lipophilic membrane stain. Additionally, SEM revealed an increase in vesicle-like structures on fungal cell surfaces. Despite these findings, the internalization of EVs by fungal cells remains an emerging field that requires further investigation to fully elucidate and consolidate our understanding of these mechanisms.

After the EV uptake, fungal cells can be triggered to mount specific responses mediated by molecules transported within these vesicles. For instance, polysaccharides like GXM found in *Cryptococcus* EVs can be assimilated by receptor cells and subsequently decorate their surface (49). Since the vesicle cargo includes polysaccharides, proteins, RNA, DNA, and other bioactive molecules, it is expected that recipient cells respond to their release within the intracellular environment. Our results revealed an increase in the mRNA expression of genes associated with virulence following co-incubation with EVs derived from the same or different fungal species. Among these genes, *Lac1* encodes laccase, an enzyme directly involved in melanin production by *Cryptococcus* spp. (50); *Ure1* encodes urease, an enzyme responsible for the hydrolysis of urea into ammonia and carbamate (51); and *Erg11* encodes sterol 14-demethylase, a key enzyme in ergosterol biosynthesis (52). The mRNA expression of all these genes was significantly altered after the addition of EVs to fungal cultures. However, further studies are needed to clarify whether the observed increase in mRNA expression is directly caused by mRNA molecules transported within EVs or if it results from other expression-activating mechanisms.

*Cap59* is a gene essential for capsule formation, as its absence or mutation results acapsular mutants (53). In our study, an increase in *Cap59* mRNA expression was conserved in cultures of ΔCap67 mutants exposed to EVs. However, TEM and India Ink staining did not reveal a restoration of the wild-type capsule phenotype. We propose two possible explanations for the reaction observed in the dot blot analysis: (a) GXM polysaccharides from EVs are being recognized as annexed structures on the cell surface and/or (b) a thin and irregularly spaced GXM layer might be produced by the acapsular mutant through the translation of *Cap59* mRNA (originating from the wild-type strain and free from missense mutations). A similar outcome was reported by Bielska et al. (2018) (23), who demonstrated that EVs from the wild-type *C. gattii* R265 strain were insufficient to restore capsule formation in an acapsular mutant (R265ΔCap10). Therefore, even though increased *Cap59* mRNA expression was detected by qPCR, this may reflect an upregulated expression of the mutated gene sequence, which would not translate into effective capsule formation.

Beyond planktonic cells, EVs can also influence fungal biofilms. Biofilm formation in fungi is a multi-stage process that includes adhesion, growth, maturation, and dispersion phases (54). Adhesion is a critical initial step, as fungal pathogens rely on this process to colonize host tissues and initiate infection. Adhesins and their corresponding host cell ligands play a pivotal role in biofilm formation (55). Research on the effects of EVs on biofilms, particularly during the adhesion phase, is still emerging. To the best of our knowledge, our study is one of the first to investigate the effects of interspecies communication in biofilm formation. While Zarnowski et al. (2022) (56) explored intra- and interspecies communication in *Candida* spp. and observed predominant antagonistic interactions between EVs from one species and biofilms from another, our findings suggest a different interaction pattern between EVs and the biofilms of *C. albicans* and *C. auris.* To address this question, we conducted experiments across a broad range of EV concentrations. Our results indicate that the concentration of EVs is a determining factor for their biological effects, whether enhancing or inhibiting biofilm adhesion, in both *Candida* and *Cryptococcus* species. Notably, EVs derived from *C. neoformans* ΔCap67 mutants did not enhance biofilm adhesion. We hypothesized that this occurs because GXM polysaccharide, which are transported by EVs from the wild-type strains, are essential for increasing adhesion levels. The absence of GXM in ΔCap67 mutants is a well-known limiting factor for biofilm formation (57).

Dispersion represents the final stage of the biofilm lifecycle, characterized by the detachment and release of cells – primarily in the yeast form - from the biofilm to colonize new surfaces (58). This process is tightly regulated by various molecules and environmental factors, including pH, nutrient availability, and other external cues. Recently, EVs are suggested as key regulators of biofilm formation and detachment (59). Previous studies using exogenous EV add-back assays in both intra- and interspecies contexts (21, 22, 58) have demonstrated that EV administration can significantly impact dispersion levels. Our findings align with these observations, showing that EVs modulated biofilm dispersion different fungal species. Interestingly, our results suggest that EVs may carry a conserved set of core effectors capable of triggering similar responses in fungi of other species. However, the biological responses elicited by EVs are not universally identical across species. Variations in response appear to correlate with the phylogenetic distance between the EV donor and recipient species, indicating that EV-mediated communication does not convey a universally consistent signal within a fungal genus (22).

In *Candida* spp., studies have linked the functional cargo transported by EVs to increased protection against antifungal treatment when EVs are added to susceptible strains (21, 22). Specifically, reduced susceptibility to fluconazole and caspofungin in *C. albicans* biofilms supplemented with EVs has been attributed to their positive role in enhancing the extracellular matrix architecture (58–60). In our study, we observed differences in antifungal tolerance between planktonic and biofilms of *C. albicans* and *C. neoformans* ΔCap67, corroborating findings from previous studies that reported a similar role for fungal EVs in both intra- and interspecies interactions (22, 40). Chan et al. (2022) (40) identified modulation of antifungal tolerance - specifically against Amphotericin B – only in interactions between fungi and EVs from the same species (*C. auris*). However, our results revealed that this modulation can also occur at the interspecies level (*C. albicans* x *C. auris*) in experiments using fluconazole. The upregulation of *Erg11* expression observed in planktonic *C. albicans* exposed to EVs, suggests that increased ergosterol synthesis could be a plausible explanation for the enhanced resistance to azoles via EV-mediated communication. Data regarding changes in antifungal tolerance following EV exposure in *Cryptococcus* spp. or other fungal pathogens, particularly in cross-species interactions, remain limited in the literature and warrant further investigation.

*Candida* spp. and *Cryptococcus* spp. are fungi belonging to distinct taxonomic classes, *Ascomycota* and *Basidiomycota*, respectively, which are phylogenetically distant (2). While *Cryptococcus* species, *Candida albicans*, and *Candida auris* are not expected to share the same ecological niches, they were utilized in this proof-of-concept study to demonstrate the principle of interspecies communication using three medically significant fungal species. Despite their genotypic and phenotypic differences, this study revealed that interspecies communication mediated by EVs can occur between evolutionary divergent fungi, resulting in biologically relevant effects on recipient cells.

Pathogenic fungal cells communicate with each other (pathogen-to-pathogen), forming a network during infection to coordinate actions through the transport of virulence factors and other molecules. They also communicate with host cells (pathogen-to-host), establishing an inter-kingdom interaction that modulates the host immune response (61). Previous studies, including work from our group, have shown that EVs from *Candida haemulonii*, *C. albicans,* and *C. neoformans* do not exhibit cytotoxic effects on macrophages. Despite this, immune cells remained capable of responding to EV stimuli through the production of cytokines, antibodies, and antimicrobial compounds, among others defense mechanisms (16, 61). Additionally, some researchers have suggested a correlation between EV size and cytotoxicity, with small EVs (sEVs) being less cytotoxic than large (lEVs) (45). Understanding the internalization of fungal EVs by innate immune cells is a rapidly evolving field. In our study, we contributed to this growing body of knowledge by demonstrating the ability of human macrophages to internalize and elicit specific responses against EVs derived from four different species of pathogenic fungi. Furthermore, our findings indicate low or absent cytotoxicity of the tested fungal EVs. This was supported by the lack of significant statistical differences in macrophage viability and cellular properties when comparing treated and untreated groups.

Pathogens invade the host and release EVs to facilitate communication between pathogenic cells, exchanging “messages” about the local environment. These messages can be intercepted by immune system cells, triggering a response against the intruder (62). The stimulation of macrophages by fungal EVs can affect cellular signaling pathways in diverse ways. This process begins with recognition by pattern recognition receptors (PRRs) and may culminate, for example, in the induction of macrophage polarization into M1 (classically activated) or M2 (alternatively activated) phenotypes (63). M1-polarized macrophages are characterized by the release of high levels of pro-inflammatory cytokines, such as IL-1β, IL-6, TNF-α, and IFNγ, and exhibit potent intracellular pathogen-killing activity mediated by inducible nitric oxide synthase (*iNOS*) and reactive oxygen species (ROS) (64). In contrast, M2 macrophages can be marked by the expression of arginase 1 (*ARG1*) and produce significant levels of anti-inflammatory, such as IL-10 and TGF-β. These cells are primarily involved in tissue repair, remodeling, and suppression of the immune response (64, 65). Arginase 1 competes with nitric oxide synthase for the substrate L-arginine, converting it into urea and ornithine. This effect contrasts with *iNOS*, which converts L-arginine into citrulline and nitric oxide (NO) (66).

Our results suggest that THP-1 macrophage-like cells were polarized toward the M1 phenotype following stimulation with fungal EVs. This pro-inflammatory behavior has also been reported for EVs from *Candida* spp. (30, 44, 67), *Cryptococcus* spp. (34), and other fungi such as *Aspergillus flavus, A. fumigatus, P. brasiliensis,* and *Tricophyton interdigitale,* among others (14, 15, 17, 68). Oliveira et al. (2010) associated stimulation of immune cells with *C. neoformans* EVs to an anti-inflammatory response. However, our findings revealed the opposite effect, a discrepancy that may be attributed to differences in the cytokines evaluated and experimental conditions. As highlighted by Brandt et al. (2024), the effects of *C. neoformans* EVs on immune cell interactions remain underexplored, underscoring a significant research gap that warrants further investigation.

While fungal recognition by immune system cells is a well-explored topic, the recognition of EVs remains underexplored. In line with observations by Honorato et al. (2024) (69), we also detected enhanced expression of *TLR4* following stimulation with *C. albicans* EVs, as well as *C. auris* EVs. This response may be linked to EV surface decorations, including *O-* and *N-* linked mannans, chitin-related structures, and β-1,3 glucans (69). TLR2 and Dectin-1 have been suggested as key receptors involved in fungal EV recognition (17, 70, 71). In our study, we observed species-specific variations in receptor expression: Dectin-1 (*CLEC7A)* was upregulated by *C. albicans* EVs, while *TLR2* was upregulated by *C. auris* and *C. neoformans* EVs. *MYD88* activation is a downstream process event in the TLR2 and TLR4 (61) signaling pathways, aligning with our findings. The detection of distinct receptor activation (or inhibition) profiles may stem from structural differences in EVs or reflect species- and strain-specific strategies employed by pathogenic fungi to evade the immune responses (61, 70). The expression of *TLR9* was a particularly noteworthy finding in our study for two reasons: (1) EVs from both *Candida* spp. significantly upregulated *TLR9,* and (2) *C. gattii* EVs significantly downregulated *TLR9*. Based on Brown et al. (2024) (3) and Vargas et al. (2020) (72), we hypothesize that the enhanced expression of *TLR9* in response to *Candida* spp. EVs may be associated with chitin and nucleic acids present in these vesicles. Conversely, the downregulation of *TLR9* by *C. gatti* EVs may represent an immune evasion strategy, as TLR9-mediated signaling cascades are crucial for host resistance against *C. gatti* infections (73).

The immunomodulatory role of Galectin-3 has previously been described in the control of *C. neoformans* and *P. brasiliensis* infections, where its upregulation was observed in infected animals. This lectin was shown to impact EV disruption, inhibit fungal growth, and enhance EV uptake by macrophages (74, 75). Our results revealed an increased expression of *LGALS3* when THP-1 cells were stimulated with fungal EVs, suggesting the involvement of this lectin not only in the recognition and destabilization of *C. neoformans* EVs but also in EVs derived from *C. albicans, C. auris,* and *C. gattii*. The recognition of *Candida* species by immune cells is typically associated with the Dectin-1 receptor and mediation through the Syk/CARD9 signaling pathway, where Syk appears to play a more direct role in antifungal immunity than CARD9 (76). However, it is also possible to observe a CARD9-dependent but Syk-independent response during *C. albicans* recognition. This mechanism involves the intracellular receptor NOD2 (*Nucleotide-binding oligomerization domain* 2) (76, 77), which may partially explain the observed increase in *CLEC7A* and *CARD9* expression alongside reduced *SYK* expression in macrophages stimulated by *C. albicans* EVs.

The contribution of STING (*Stimulator of Interferon Genes*) during fungal infections remains under discussion, as its activation has been shown to play a negative modulatory role in antifungal immunity following recognition of fungal EVs (78–80). Harding et al. (80) used *C. albicans* EVs to stimulate bone marrow-derived macrophages for 6 hours, observing activation of the cGAS (*cyclic GMP-AMP synthase*)-STING pathway. However, our results demonstrated *STING* mRNA downregulation in macrophages stimulated with all fungal EVs tested. We propose that this discrepancy may stem from several factors, including differences in fungal strains, vesicle cargo composition (which depends on fungal culture conditions), cell lineages, macrophage stimulation duration (6 h vs. 20 h), immunomodulatory effects of fungal EVs, or even a macrophages strategy to mitigate excessive STING activation, which could be detrimental to immune cells. In summary, immune responses and activated pathways are both fungal- and EV composition-dependent (81). In addition to STING, CGAS is another key component of this pathway. Its downregulation has been linked to increased levels of pro-inflammatory cytokines (*IL-1β, IL-6,* and *TNF-α*) in the context of *A. fumigatus* infections (82). Following THP-1 stimulation with EVs, *C. neoformans* – and especially *C. gattii* - EVs induced *CGAS* downregulation. This suppression mechanism may negatively impact the immune response, facilitating fungal immune evasion, as suggested by Peng et al. (2023) and Gu et al. (2023) (78, 82).

The characterization of immune response following exposure of host innate immune cells to fungal EVs has advanced through cytokine secretion analysis, enabling the identification of patterns suggestive of pro- or anti-inflammatory responses (62). EVs from *C. neoformans, C. gattii, C. albicans,* and *C. auris* were previously described as to elicit pro-inflammatory effects in immune cells (32). These responses typically involved increased production of TNFα, IL-6, IL-8, and IL-12, alongside reduced or less pronounced production of TGF-β and IL-10 (61). Consistent with previously published data on fungal EV immunomodulation, our results indicated a predominance of pro-inflammatory cytokines (IL-1β, IL-6, IL-8, and TNFα) in EV-treated macrophages, accompanied by reduced IL-4 levels. Similarly, decreased anti-inflammatory cytokine production compared to non-treated macrophages has been reported in other studies (44, 67, 83). Despite the overall pro-inflammatory profile, IL-10 secretion was observed at lower levels. A pro-inflammatory response following EV recognition may represent a preparatory mechanism to combat fungal infections (62). This inter-kingdom cross-communication between host defense mechanisms and pathogenic fungi highlights a crucial area for further exploration, potentially guiding novel interventions for patients affected by invasive fungal infections (84).

Taken together, the results obtained in this study represent valuable tools for the development of antifungal therapies. We demonstrated that fungal cells can communicate with each other via signals carried by EVs, which are interpreted by receptor cells from the same or different species. Although the fungal species studied here are unlikely to be found together in the environment or during infection, they belong to fungal genus that include major human pathogenic fungi as thus serve as representative examples to explore EV-mediated interspecies communication. Using *C. albicans, C. auris, C. neoformans,* and *C. gattii,* as models pathogenic fungi, along with their respective EVs, we showed that these molecular messages can also be directed to or intercepted by immune system cells, exerting significant effects on various targets of the innate immune response. Therefore, investigating fungal EVs offers critical insights into fungal communication network, highlighting the need for further evidence to fully “decode” the mechanisms underlying fungal orchestration during infections.

## Materials and methods

### Strains and culture conditions

*Candida albicans* ATCC 64548 and a clinical isolate of *Candida auris* 470/2015 (clade IV) were cultured in Sabouraud liquid or solid medium (2% w/v agar) at 37 °C for 48 h, with or without shacking at 200 rpm. *Cryptococcus neoformans* var. *grubii* H99, *Cryptococcocus gattii* R265 (kindly supplied by Prof. Dr. Thiago Aparecido da Silva, UNESP), and the *Cryptococcus neoformans* var. *neoformans* Cap67 mutant (ΔCap67) were cultured in a chemically defined minimal medium containing 15 mM dextrose, 10 mM MgSO_4_, 29.4 mM KH_2_PO_4_, 13 mM glycine, and 3 μM thiamine. These cultures were incubated under the same conditions described above, as minimal media are associated with increased EVs production (85). The ΔCap67 mutant carries a missense mutation in the *Cap59* gene (position 1345), resulting in an amino acid substitution from glycine (Gly) to serine (Ser). This mutation eliminates capsule formation, resulting in an acapsular strain (Cap^-^ phenotype) (36, 53).

The human monocytic cell line THP-1 (ATCC TIB202) was cultured in RPMI 1640 medium (Gibco®) supplemented with 10% (v/v) exosome-depleted fetal bovine serum (FBS) (Vitrocell®), sodium bicarbonate, and penicillin/streptomycin (1000 IU). Cells were maintained in a humidified incubator at 37 °C with 5% CO_2_. Differentiation of THP-1 monocytes into adherent macrophage-like cells was induced by treating the cultures with 10 ng/mL Phorbol 12-myristate 12-acetate (PMA) for 24 h.

### Isolation of EVs

EVs from all yeasts strains used in this study were isolated following the protocol described by Reis et al. (2019) (39). Briefly, overnight cultures of each yeast strain in 10 mL of Sabouraud medium were counted using a Neubauer chamber. A total of 3,5 × 10^7^ cells were inoculated onto each 2% agar Sabouraud plate, with 10 plates per strain used to ensure sufficient EV yield. After 48 h of incubation at 37 °C, yeast material was recovered from the plates using Phosphate-Buffered Saline (PBS). The eluate was centrifuged in a Sorvall LEGEND RT+ (Thermo®) centrifuge at 5,000 × *g* for 15 min to remove larger debris. The resulting supernatant was further centrifuged at 15,000 × *g* for 15 min to eliminate residual cellular material, and the pelleted cells were counted to determine the EV-to-cell ratio. The supernatant was passed through 0.45 μM membrane filters (CORNING®) to remove any remaining debris. Finally, EVs were pelleted by ultracentrifugation at 100,000 × *g* for 1 h at 4 °C using an Optima MAX-XP ultracentrifuge (Beckman®). The EV-enriched pellet was resuspended in ultrapure water (Sigma-Aldrich®) and stored at – 80°C. Subsequent EV analyses were performed in accordance with the MISEV 2023 guidelines (*Mininal Information for Studies of Extracellular Vesicles*) (12).

### Nanoparticle tracking analysis (NTA)

Nanoparticle tracking analysis was conducted to characterize the size distribution and quantify fungal EVs. Analysis was performed on a Nanosight NS300 (Malvern®) equipped with NTA 3.0 software, following the manufacturer’s instructions. Samples were diluted at a 1:100 ratio in PBS and carefully loaded into the Nanosight system. System parameters were adjusted to detect 20–100 particles per frame, with camera sensitivity set to 14 or above (16). Results of NTA were plotted using OriginPro® 2024 software (OriginLab ©).

### Zeta potential

The zeta potential of fungal EVs was measured to evaluate surface charge and colloidal stability, using laser scattering with a Zetasizer Nano ZS (Malvern®) (λ = 580 nm, scattering angle 172°), equipped with Zetasizer software v. 7.1 (Malvern®). Measurements were performed at 25 °C in a disposable DTS1070 cuvette using PBS as the dispersant (pH 7.4). Electrophoretic mobility values were converted to zeta potential using the Smoluchowski equation, and each sample was analyzed in triplicate with 10– 100 runs per measurement.

### Transmission Electron Microscopy (TEM)

Fungal EV samples (1 × 10^9^ EVs) were fixed in a solution containing 2% glutaraldehyde and 2% paraformaldehyde in 0.1 M sodium cacodylate buffer (pH 7.4) for 4 h at 4 °C. Following fixation, samples were ultracentrifuged at 100.000 × g at 4 °C for 1 h, and the resulting pellets were resuspended in 200 μL of sodium cacodylate buffer. EV structures were visualized using a JEM 100CXII transmission electron microscope (JEOL®), and images were acquired with a Hamamatsu ORCA-HR digital camera (JEOL®).

### Internalization of EVs by fungal cells and human macrophages

The recognition and internalization of EVs secreted by either the same or different fungal species were analyzed using fluorescence microscopy. A volume corresponding to 1 × 10^9^ EVs was stained with 1 uL of Vybrant Dil Cell-Labeling Solution (5 μM, Invitrogen®), a lipophilic fluorescent dye for membrane labeling, and incubated for 20 min at 37 °C. After staining, excess dye was removed by washing with PBS. The samples were subsequently ultracentrifuged at 100.000 × *g* at 4 °C for 1 h, and the resulting EV pellet was resuspended in Sabouraud liquid medium, previously inoculated with fungal cells at a 1000:1 EV-to-fungus ratio, and incubated under agitation at 200 rpm.

### Scanning Electron Microscopy (SEM)

Scanning electron microscopy was employed to visualize fungal cell morphology following incubation of *C. albicans* and *C. neoformans* (ΔCap67 mutant) with EVs in Sabouraud medium at a concentration of 1 × 10^9^ EVs and 1 × 10^6^ cells (1000:1 EV-to-fungus ratio) under the following conditions: a) *C. albicans* with EVs from *C. auris*, and b) *C. neoformans* ΔCap67 with EVs from *C. gattii*. Incubation was performed at 37 °C with agitation at 200 rpm over 24 h. After incubation, fungal cells were centrifuged at 5,000 × g for 15 min and washed with PBS to remove unbound EVs. Cells were then fixed with 2% paraformaldehyde diluted in PBS for 2 h at 4 °C, centrifuged again as previously described, and resuspended in a buffer containing 1% osmium tetroxide (OsO_4_). Samples underwent dehydrated through a graded ethanol series, followed by critical point drying using an Autosamdri-810 system (Tousimis®). Dried samples were mounted on aluminum stubs with silver paint and coated with gold using a Sputter Coater SCD 050 (Bal-Tec). Finally, the prepared samples were visualized using a JSM-6610 LV scanning electron microscope (JEOL®).

### Dot blot

The transfer of virulence factors via EVs was investigated by evaluating the transmission of GXM, the primary component of the cryptococcal capsule, to a non-capsular mutant strain of *C. neoformans* (ΔCap67). A straightforward and effective assay was employed, wherein *C. neoformans* ΔCap67 was incubated with EVs from *C. neoformans* and *C. gattii.* In this assay, *C. neoformans* ΔCap67 cells (1 × 10^6^/mL) were incubated with EVs (1 × 10^9^/mL) derived from *C. neoformans* and *C. gattii* in Sabouraud medium for 24 h at 37 °C with 5% CO_2_. After incubation, the cells were harvested by centrifugation at 8.000 × *g* for 15 min, washed with PBS, and 10 μL of each sample were applied to a Polyvinylidene difluoride (PVDF) membrane pre-treated with methanol for 5 min. The membrane was subsequently blocked with 5% (w/v) skim milk in PBS for 30 min and washed three times with PBS-T (PBS + 0.05% Tween 20). Next, the membrane was incubated with the monoclonal antibody 18b7, which specifically binds to GXM (generously provided by Prof. Dr. Anamélia Bocca from FIOCRUZ), for 1 h. Following another washing step, an anti-mouse HRP-conjugated secondary antibody (Sigma®) was applied for 1 h. After five additional washes with PBS, the membrane was treated with a detection solution (Vector Laboratories®), resulting in a visible signal indicating the presence of GXM in the samples.

### Co-incubation of *C. albicans* and acapsular *C. neoformans* planktonic cells with EVs

*C. neoformans* ΔCap67 were grown to the stationary phase in Sabouraud broth. A volume corresponding to 5 × 10^6^ cells from the previous culture was inoculated into 5 mL of fresh Sabouraud broth and mixed with 1 × 10^9^ EVs (1:200 cell/EV ratio) isolated from *C. neoformans* H99, *C. gattii* R265, or self-derived EVs from ΔCap67. Fungal EVs were added at the start of the experiment (0 h) and again after 6 h of incubation. The experiment was conducted over 24 h at 37 °C with shaking at 200 rpm. At designated time points, samples were collected, and India ink staining was performed to visualize potential capsule formation on the cellular surface. Cells were then separated from the culture medium by centrifugation at 8.000 × *g* for 15 min at 4 °C and subjected to RNA extraction using the Quick-RNA Fungal/Bacterial Miniprep kit (Zymo Research®). The isolated RNA was quantified using a NanoDrop One spectrophotometer (Thermo®) and subsequently used to evaluate the expression levels of *Lac1*, *Ure1,* and *Cap59* by qPCR, with results compared to the control culture (*C. neoformans* ΔCap67 without EVs addition).

The effect of fungal EVs on planktonic cells of *Candida* was investigated using a similar experimental design. Cultures of *C. albicans* were first grown to the stationary phase in Sabouraud broth. The same broth volume, cell/EV ratio, and incubation conditions previously described were applied. In this assay, EVs isolated from *C. albicans* and *C. auris* were analyzed. Additionally, EVs from *C. auris* cultured in the presence of fluconazole were included for comparison. After incubation, the cell pellet was collected by centrifugation, and total RNA was purified and quantified as described before for the *C. neoformans* ΔCap67 experiment. The expression level of *Erg11* was then evaluated by qPCR and compared to a control culture (*C. albicans* without EV treatment).

### Spot test

The effect of GXM-transmitting EVs on *C. neoformans* ΔCap67 cells was further investigated using a spot assay to assess potential differences in antifungal tolerance. In this assay, 100 μL of EVs (1 × 10^10^ EVs/mL) derived from *C. neoformans* or *C. gattii* were added to cultures of *C. neoformans* ΔCap67 (1 × 10^6^ cells/mL) in 10 mL of Sabouraud broth. The cultures were then incubated at 37 °C with shaking at 200 rpm for 24 h. Untreated *C. neoformans* ΔCap67 cultures served as controls. Following incubation, the cultures were centrifuged at 8.000 × *g* for 15 min, washed with PBS, and subjected to tenfold serial dilutions, starting from 10^7^ cells/mL in PBS. Subsequently, 10 μL of each dilution was plated on Sabouraud agar (control) and Sabouraud agar supplemented with fluconazole (1 - 6 μg/mL). Plates were incubated at 37 °C for 96 h.

To further explore the effect of EVs on antifungal drug tolerance, a similar spot assay was performed using *C. albicans* (fluconazole-susceptible) cultures (1 × 10^6^ cells/mL). These cultures were mixed with 100 μL of EVs (1 × 10^10^ EVs/mL) derived from a fluconazole-resistant *C. auris* strain and incubated in 10 mL of Sabouraud broth at 37 °C with shaking at 200 rpm for 24 h. Control cultures of *C. albicans* and *C. auris* without EV treatment were also included. Following incubation, the cultures were centrifuged, washed, and serially diluted tenfold. 10 μL of each dilution were plated on Sabouraud agar and Sabouraud agar supplemented with fluconazole (1000 μg/mL). Plates were incubated at 37 °C for 96 h.

### Biofilm adhesion

The effect of isolated EVs on biofilm adhesion was evaluated following the protocol described by Zarnowski et al. (2021) (21), using a 96-well microplate assay. Each well was inoculated with 1 × 10^6^ cells of *C. albicans* or *C. auris* in RPMI medium. Different concentrations of EVs (10-fold dilutions ranging from 10^9^ to 10^5^) were added to the culture medium to assess their impact on biofilm formation. After 90 min of incubation at 37 °C with 5% CO_2_, the culture medium was carefully removed to eliminate non-adherent cells. The biofilms were then washed with 100 μL of PBS. Biofilms adhesion was quantified using the MTT (*Dimethyl-2-thiazol tetrazolium bromide*) assay. For this, MTT was prepared in PBS at a final concentration of 0.5 mg/mL and protected from light. A volume of 100 μL of this MTT solution was added to each well, and the plate was incubated for 3 h at 37 °C. Cellular dehydrogenase activity in living cells can be followed by the reduction of MTT to formazan, a highly colored compound, due to the action of NADH. The formazan crystals were then dissolved in 100 μL of DMSO (*dimethyl sulfoxide*) was added, followed by an additional 10-min incubation at 37 °C with gentle shaking at 50 rpm. Absorbance was measured at 490 nm using a SpectraMax 190 Microplate Reader spectrophotometer (Molecular Devices®).

The effect of EVs on biofilm adhesion of *C. neoformans* ΔCap67 was also evaluated based on protocols by Martinez et al. (2006) (57) and Benaducci (2016) (51), with modifications. Briefly, *C. neoformans* ΔCap67 was cultured overnight in Sabouraud broth at 37 °C with shaking at 200 rpm. Cells were then counted using a Neubauer chamber, centrifuged at 8.000 × *g* for 15 min, and resuspended in minimal medium (20 mg/mL thiamine, 30 mM glucose, 26 mM glycine, 20 mM MgSO_4_ ·7H_2_O, 58.8 mM KH_2_PO_4_) to a final concentration of 1 × 10^7^ cells/mL. In a 96-well microplate, 2 × 10^6^ cells were added to each well and mixed with EVs at different concentrations (10-fold dilutions ranging from 10^9^ to 10^5^) in a final volume of 200 μL. After the adhesion stage, the supernatant was carefully aspirated and discarded, and biofilms were washed with PBS to remove non-adherent fungal cells. Biofilm adhesion was subsequently assessed using the MTT assay, following the procedure described above.

### Biofilm dispersion

The impact of EVs on preformed *Candida* biofilms was assessed using a dispersion assay, following adapted protocols from Zarnowski et al. (2021) (21) and Zarnowski et al. (2022) (22). The assay was performed in 96-well microplates, with each well inoculated with 100 μL of a *C. albicans* or *C. auris* suspension (1 × 10^7^ cells/mL) in RPMI medium. After a 90-min incubation at 37 °C with 5% CO_2_, the culture medium was aspirated, and the biofilms were washed with 100 μL of PBS. Fresh RPMI medium was then added, and the biofilms were incubated for an additional 24 h under the same conditions. EVs isolated from *C. albicans* and *C. auris* were prepared at concentration ranging from 1 × 10^9^ to 1 × 10^7^ EVs/mL in RPMI medium - concentrations previously shown to affect the adhesion stage. These EV solutions were added to the preformed biofilms, followed by a 24-hour incubation. Afterward, supernatants containing detached cells were carefully transferred to a new microplate, where the quantity of dispersed cells was assessed using an MTT assay with the reagent at double concentration. Biofilm dispersion was analyzed by comparing EV-treated biofilms with untreated control biofilms.

The dispersion stage, characterized by the detachment of microcolonies or planktonic cells from mature *Cryptococcus* biofilms (86), was also evaluated in response to cryptococcal EVs. Following protocols adapted from studies on *Candida* biofilms (21, 22) and *Cryptococcus* biofilms (51, 57), a dispersion assay was performed in 96-well microplates. Each well was inoculated with 2 × 10^6^ cells of *C. neoformans* ΔCap67 in minimal medium. After the initial adhesion stage, biofilms were washed with PBS to remove non-adherent cells and allowed to mature at 37 °C for 48 h in RPMI medium. The medium was then carefully replaced with suspensions containing 1 × 10^9^ EVs from *C. neoformans* ΔCap67, *C. neoformans* H99 or *C. gattii* R265, diluted in RPMI. Following an additional 24-hour incubation, biofilm dispersion was quantified using the same MTT assay protocol applied to *Candida* biofilms.

### Antifungal Susceptibility of Biofilms

Biofilm susceptibility to antifungal treatment was evaluated using a vesicle add-back assay, following the protocols described by Karkowska-Kuleta et al. (2023) (58) and Zarnowski et al. (2021) (21). Biofilms of *C. albicans* were established by inoculating 1

× 10^5^ cells per well in a 96-well microplate. After a 90-min incubation at 37 °C in a humidified atmosphere with 5% CO_2_ to allow initial adhesion, the culture medium was aspirated, and biofilms were washed with PBS. Subsequently, 100 μL of RPMI medium containing EVs at concentrations of 1 × 10^9^ EVs/mL were added to the wells, followed by a 2-hour incubation at 37 °C. Fluconazole was then diluted in RPMI to a final concentration of 100 μg/mL and added to the wells, bringing the total volume to 200 μL per well. Control wells included biofilms without EV treatment. The microplates were incubated for 24 h at 37 °C with 5% CO_2_ in a humidified environment without agitation. Biofilm susceptibility to fluconazole was assessed using the MTT assay.

The influence of EVs on cell communication and antifungal susceptibility in *C. neoformans* ΔCap67 was evaluated using a microplate assay adapted from Martinez et al. (2006) (57). *C. neoformans* ΔCap67 cells were cultured in Sabouraud liquid medium for 48 h at 37 °C with shaking at 200 rpm. The cultures were centrifuged at 8.000 × *g* for 15 min, and the resulting pellet was washed with PBS, centrifuged again, and resuspended in RPMI medium to a final concentration of 1 × 10^7^ cells/mL. In a 96-well flat-bottom microplate, each well received 100 μL of the *C. neoformans* ΔCap67 suspension (1 × 10^7^ cells/mL), 100 μL of EVs (1 × 10^9^ EV/mL diluted in RPMI) derived from either *C. neoformans* or *C. gattii,* and 100 μL of antifungal agents (fluconazole at 10 μg/mL or amphotericin B at 5 μg/mL, both diluted in RPMI). The microplates were incubated at 37 °C with shaking at 50 rpm for 72 h. Absorbance was measured at 600 nm (OD _600nm_) using a SpectraMax 190 Microplate Reader (Molecular Devices®).

### Stimulation Assay of THP-1 macrophages with fungal EVs

For the stimulation assay, 6-well plates were prepared with 1 × 10^6^ THP-1 cells per well, performed in triplicate. EVs were diluted in RPMI medium to a final concentration of 1 × 10^9^ EVs/mL, and 1 mL of this solution was added to each well, resulting in an EV-to-macrophage ratio of 1000:1. Lipopolysaccharide (LPS) (Sigma®) at 100 ng/mL was included as a positive control, while RPMI medium without EVs served as the negative control. The plates were incubated for 20 h at 37 °C with 5% CO_2_. Following incubation, supernatants were carefully collected and stored at –20 °C for subsequent cytokine quantification. Cells were harvested using PBS, and total RNA was extracted using the RNeasy Mini kit (Qiagen®) according to the manufacturer’s protocol. RNA concentration and purity were determined using a Nanodrop One spectrophotometer (Thermo®), and the extracted RNA was stored at -80 °C for later qPCR analysis. To evaluate the impact of EVs on THP-1 cells, cell size, viability, and concentration were assessed using the Countess 3.0 Automated Cell Counter (Thermo®).

### Detection of Cytokine Production

Culture supernatants were pooled according to the respective stimulus and stored at –20 °C until cytokine quantification via enzyme-linked immunosorbent assay (ELISA). Human cytokine detection kits for TNFα, IL1β, IL4, IL6, IL8, and IL10 (BD Biosciences®) were used following the manufacturer’s instructions. Cytokine levels were determined by measuring absorbance at 450 nm, with wavelength correction at 570 nm, using a SpectraMax 190 Microplate Reader (Molecular Devices®).

### Quantitative Real-time PCR (qPCR)

For cDNA synthesis, 1 μg of total RNA obtained from the THP-1 stimulation assay was reverse-transcribed using the High-capacity cDNA Reverse Transcription Kit (Applied Biosystems®), following the manufacturer’s protocol, including the recommended time and temperature conditions. Quantitative PCR (qPCR) was performed using the SyGreen Mix (PCR Biosystems®) in a final reaction volume of 10 μL per well, containing 50 ng of cDNA. Reactions were conducted on a StepOne Plus Real-time PCR system (Applied Biosystems®) under the following cycling conditions: initial denaturation at 95 °C for 2 min, followed by 40 cycles of 5 sec at 95 °C and 30 sec at 60 °C, as per PCR Biosystems ® recommendations. In the fungal communications experiments (fungal culture + EVs), the mRNA expression levels of *Erg11*, *Lac1, Ure1,* and *Cap59* were normalized to the housekeeping gene *18S rRNA*. Additionally, the immune response triggered by fungal EVs in THP-1 macrophage cultures was evaluated via qPCR by analyzing the expression of *TLR2, TLR4, TLR9, LGALS3* (Galectin-3)*, CLEC7A,* (Dectin-1), *ARG1, iNOS*, *CGAS, STING, SYK, MYD88*, and *CARD9* (refer to Supplementary Table 1 for primers list). These genes were normalized against the housekeeping gene *ACT1* (β-Actin). Relative mRNA expression levels were calculated using the 2-ΔΔCT method, as described previously (87).

### Statistical Analysis

Data analysis was performed using GraphPad Prism 9.0 (Dotmatics®). Statistical comparisons were conducted using either one-way or two-way ANOVA, depending on the experimental design. A p-value of p < 0.05 was considered statistically significant.

## Acknowledgments

The authors thank Carlos Alberto Vieira, Maria Dolores Seabra Ferreira, and José Augusto Maulin for technical support. This study was financed, in part, by the São Paulo Research Foundation (FAPESP), Brazil. Process Number 2023/05800-7; 2021/06794-5.

